# An autonomous, but INSIG-modulated, role for the Sterol Sensing Domain in mallostery-regulated ERAD of yeast HMG-CoA reductase

**DOI:** 10.1101/2020.08.20.260133

**Authors:** Margaret A Wangeline, Randolph Y Hampton

**Affiliations:** From the Division of Biological Sciences, the Section of Cell and Developmental Biology, UCSD, La Jolla, California 92093

**Author notes:** Corresponding author: Randy Hampton, University of California San Diego, 9500 Gilman Dr., La Jolla, CA 92093-0347.

**Keywords:** Endoplasmic-reticulum-associated protein degradation (ERAD), ER quality control, protein misfolding, cholesterol regulation, ubiquitin, ubiquitin-proteasome system, HRD pathway, HMG-CoA Reductase, ssterol sensing domain (SCAP), mallostery

## Abstract

HMG-CoA reductase (HMGR) undergoes feedback regulated degradation as part of sterol pathway control. Degradation of the yeast HMGR isozyme Hmg2 is controlled by the sterol pathway intermediate GGPP, which causes misfolding of Hmg2 to enhance its ERAD by the HRD pathway. GGPP-dependent reversible misfolding of Hmg2 is remarkably similar to classic allosteric control; we recently labeled this process mallostery to fuse the ideas of misfolding and allostery. We have evaluated the role of the Hmg2 sterol sensing domain (SSD) in mallostery, and the involvement of highly conserved INSIG proteins in SSD function. The SSD is a membrane-embedded motif found in many sterol-related proteins. The Hmg2 SSD was critical for in vivo regulated degradation of Hmg2, and required for mallosteric misfolding of GGPP as studied by in vitro limited proteolysis. The Hmg2 SSD functions in mallostery independently of conserved yeast INSIG proteins. However, this autonomous action of the SSD was modulated by INSIG, thus imposing a second layer of control on Hmg2 regulation. SSD-mediated mallostery occurs prior to HRD dependent ubiquitination, defining a pathway regulation involving SSD-mediated misfolding followed by HRD dependent ubiquitination. GGPP dependent misfolding occurred at a much slower rate in the absence of a functional SSD, indicating that the SSD functions to allow physiologically useful rate of GGPP response, and implying that the SSD is not a binding site for GGPP. We used unresponsive Hmg2 SSD mutants to test the importance of quaternary structure in mallosteric regulation: the presence of a non-responsive Hmg2 mutant strongly suppressed regulation of a co-expressed, normal Hmg2. Finally, we have found that GGPP regulated misfolding occurred in detergent solubilized Hmg2, indicating that the mallosteric response is an intrinsic feature of the Hmg2 multimer. The preserved response of Hmg2 when in micellar solution will allow next-level studies on the structural and biophysical features of this novel fusion of regulation and protein quality control.

## Introduction

ER-associated degradation (ERAD) refers to a conserved set of degradation pathways that detect and degrade misfolded, unassembled, and damaged ER-resident proteins (1–4). Both lumenal (ERAD-L) and integral membrane (ERAD-M) proteins can be subject to ERAD. ERAD is initiated by the action of a surprisingly small set of conserved E3 ubiquitin ligases that each recognize a large range of substrates. The two major ERAD pathways in yeast—defined by the participant ligases—are the HRD (pronounced “herd”) pathway and the DOA (pronounced “dee-oh-ay”) pathway. The detailed structural features of an ERAD substrate that determine HRD- or DOA-dependent degradation are being unraveled, and appear to encompass a large number of variations from wild-type stable configuration. Despite the large variety of possible substrates accommodated by each pathway, ERAD displays high specificity for misfolded versions of the proteins that undergo degradation, as would be required for an evolutionarily successful quality control pathway.

In the course of our studies on the sterol synthetic pathway in yeast, we discovered that the HRD ERAD quality control pathway is also used to regulate levels of the normal, rate-limiting sterol synthetic enzyme HMG-CoA reductase (HMGR or HMGcR) (5–7). Specifically, the Hmg2 isozyme undergoes negative feedback regulation effected at the level of HRD-dependent degradation: the 20-carbon sterol pathway molecule geranylgeranyl pyrophosphate (GGPP) accelerates HRD-dependent degradation of Hmg2, thus allowing control over Hmg2 levels keyed to changing cellular demand for sterol pathway products. Interestingly, although the regulatory mechanisms are distinct, mammalian HMGcR stability is also controlled in part by GGPP-mediated enhancement of degradation by ER-localized E3 ligases (8).

Like other quality control pathways, HRD-mediated ERAD is highly specific for misfolded versions of its many protein substrates. Our studies have revealed that the high selectivity of the HRD pathway for misfolded versions of substrates underlies HRD-dependent regulation of Hmg2 stability by GGPP: when GGPP levels are elevated, Hmg2 undergoes a reversible structural transition to a more misfolded form, allowing enhanced recognition and destruction by the HRD pathway and consequent lowering of Hmg2 activity. It appears that this mode of regulation by quality control has many examples throughout biology(9–16). Furthermore, the presence of numerous degradative quality control pathways in all cellular compartments allows for the possibility of translational applications of regulated quality control in which small molecules that program quality control degradation of desired targets could be discovered and developed. Accordingly, we have devoted significant energy towards understanding the mechanisms at play in the GGPP-mediated enhancement of HRD dependent Hmg2 degradation.

We have employed a variety of approaches to observe that GGPP causes reversible misfolding of the Hmg2 protein to enhance HRD-dependent Hmg2 degradation (17–19). Using both in vivo and in-vitro methods, we found that the effects of GGPP on Hmg2 structure are remarkably similar to allosteric regulation. Because of these similarities, we have named this ligand-based misfolding “mallostery”, a portmanteau combining the ideas of misfolding with those of allosteric regulation to refer to this type of highly specific ligand mediated misfolding (13, 18).

In our studies of mallostery, we built the case for the allosteric analogy in Hmg2 regulated degradation, showing high potency and structural specificity of GGPP, evidence of specific binding indicated by a close analog -GGSPP-that functions as a GGPP antagonist, and that reversible misfolding is central to GGPP’s role as an indicator of sterol pathway activity. In the studies below, we examined the in-cis features of Hmg2 regulation by GGPP, using a variety of Hmg2 mutants with highly specific lesions in degradative behavior to both explore the nature of ligand-mediated misfolding, and to further test the mallosteric model. We paid particular attention to the highly conserved sterol sensing (SSD) domain, found in many proteins that pertain to sterol response, synthesis, or transport. The data indicated that the SSD functions as an autonomous motif to promote mallosteric misfolding. Furthermore, the mutants described in these studies allowed a strong test of the importance of the multimeric structure in ligand-regulated Hmg2 misfolding. Finally, we demonstrate that mallosteric regulation of Hmg2 can occur in detergent solubilized Hmg2, setting the stage for next level experiments to both understand and harness this highly specific regulatory strategy in both fundamental and translational endeavors.

## Results

### The sterol-sensing domain (SSD) is required for ligand-regulated Hmg2 misfolding

The sterol-sensing domain (SSD) is a motif found in the multispanning membrane domains of many proteins that function in sterol synthesis, response, or regulation, including yeast Hmg2 (13, 19–22). This highly conserved motif occurs in the intra- and juxta-membrane residues over five adjacent membrane-spanning alpha helices. Often, mutation of highly conserved SSD residues alters the regulatory or functional responses of SSD-containing proteins (19, 23, 24). Our previous work identified several conserved residues in the SSD of Hmg2 that are required for Hmg2 regulated degradation (Fig. 1A) (19). Surprisingly, all SSD mutants that caused changes in Hmg2 degradation, including mutation of the highly conserved S215 residue to A produced *increased* stability over wild-type (19), in contrast to the usual role of strongly conserved residues in permitting maximal stability. We had earlier noted that the S215A mutation also blocked the response to the putative regulatory ligand farnesol (FOH) in an *in vitro* limited proteolysis assay of Hmg2 structure. Later work demonstrated that the *bona fide* signal for Hmg2 regulated degradation is the highly potent, mallosteric regulator geranylgeranyl pyrophosphate (GGPP), which is effective at concentrations approximately 1000 times lower than FOH (18). Accordingly, we examined the importance of these phenotypic, conserved SSD mutations on physiologically relevant, GGPP-induced reversible misfolding of Hmg2, using both in vivo and in vitro approaches.

**Figure 1.**
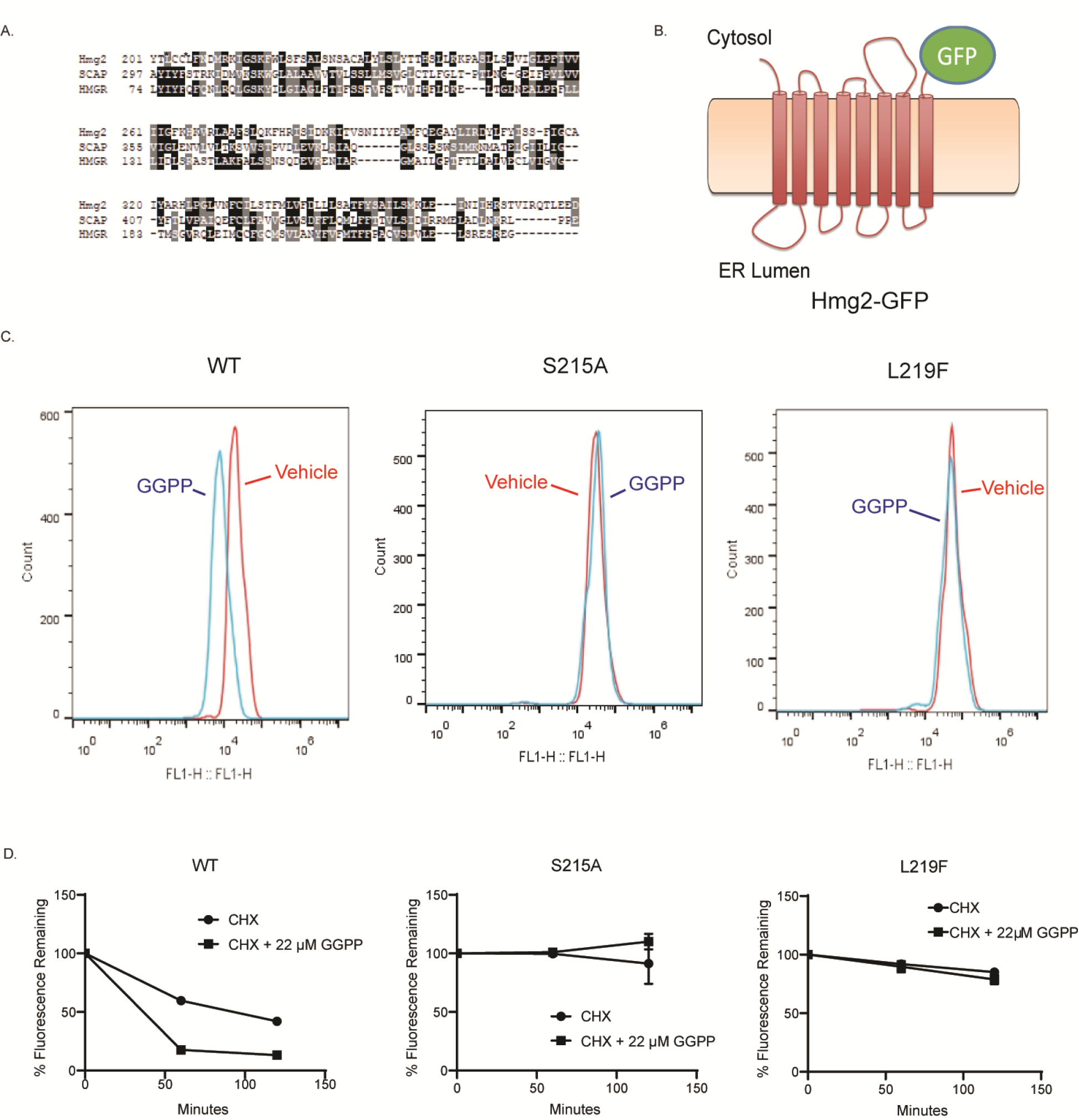
The Hmg2 sterol sensing domain (SSD) was required for GGPP-regulated degradation A. Amino acid sequence alignment of S. cerevisiae Hmg2 and H. sapiens SCAP and HMGR containing the region of the SSD. Similarities are highlighted in gray and identities are highlighted in black. Asterisks show the conserved residues S215 and L219. B. Schematic of the optical reporter Hmg2-GFP. The transmembrane region of Hmg2 is fused to GFP. (Cartoon originally appeared in (18) and used with permission.) C. SSD mutations rendered Hmg2 more stable in vivo. Histograms of Hmg2-GFP, wild-type or with the mutations S215A and L219F, as indicated, at steady state (red) or after two hours of cycloheximide treatment (blue). D. Stabilized SSD mutations are GGPP insensitive. Cycloheximide chase of wild-type and the S215A or L219F SSD mutants of Hmg2-GFP, in which CHX is added at time 0 and subsequent levels were measured by flow cytometry, in the presence of vehicle (solid circles) or 22μM GGPP (solid squares). Error bars are SEM.

To study the role of the Hmg2 SSD in vivo and in vitro, we employed the Hmg2-GFP reporter, in which the C-terminal catalytic domain of Hmg2 is replaced by GFP (Fig. 1B). Hmg2-GFP lacks HMG-CoA reductase catalytic activity but undergoes quantitatively normal regulated degradation (18, 25, 26). We focused on the mutants with the strongest phenotype from our earlier broad survey of this motif. The two mutants featured herein, S215A and L219F, each showed a loss of response to direct addition of the degradation signal GGPP to the living cells. Both mutants were non-responsive GGPP addition compared to wild-type Hmg2-GFP (Fig. 1C). Similarly, both mutants displayed the expected strong stabilization after addition of cycloheximide, while the wild-type protein displayed a drop in steady state levels due to degradation stimulated by the natural ambient concentration of GGPP (Fig 1D; (19); this lack of degradation in response to GGPP can be further assayed by direct addition of GGPP to during cycloheximide chase to stimulate HRD-dependent Hmg2 degradation (18, 26); Again, the two mutants were essentially nonresponsive to GGPP addition (Fig 1D). This confirmed that the GGPP stimulated degradation showed the expected dependence of highly conserved SSD residues predicted from our earlier studies, allowing us to delve more deeply into the role of this motif in mallosteric control of Hmg2 folding.

To that end, we employed a limited proteolysis assay of Hmg2, involving a myc-tagged version of Hmg2-GFP called (1myc_L_-Hmg2-GFP)-developed early in our studies of regulated degradation-in our analysis of the SSD. The myc tag in 1myc_L_-Hmg2-GFP is present in the first luminal loop of Hmg2 and so is protected by the ER membrane, allowing detection of the protein and its cleavage products after proteolytic treatment of the Hmg2 in isolated microsomes (Fig. 2A). Importantly, 1myc_L_-Hmg2-GFP is regulated normally (18, 27, 28). When microsomes prepared from strains expressing 1myc_L_-Hmg2-GFP are treated with limiting concentrations of trypsin, a characteristic pattern of cleavage fragments is produced that can be detected by blotting for the protected myc tag, and example being shown in 2A. When GGPP is included in the proteolysis assay, Hmg2 proteolysis occurs more rapidly ((18, 28), and Fig2B, C left sides of panels). Only the rate at which the fragments are produced changes but not the pattern itself (27, 28). This assay was used extensively in our in initial characterization of GGPP-mediated mallostery (18). The highly stable S215A version of 1myc_L_-Hmg2-GFP did not respond to added GGPP at any concentration tested, even 2000-fold higher than that required to stimulate misfolding in the wild type protein (Fig 2C). Similarly, the stabilizing L219F mutant of the conserved SSD residue also blocked the response to GGPP in vitro (Fig. 2D). Thus, it appears that GGPP dependent misfolding occurred by an SSD-dependent process.

**Figure 2.**
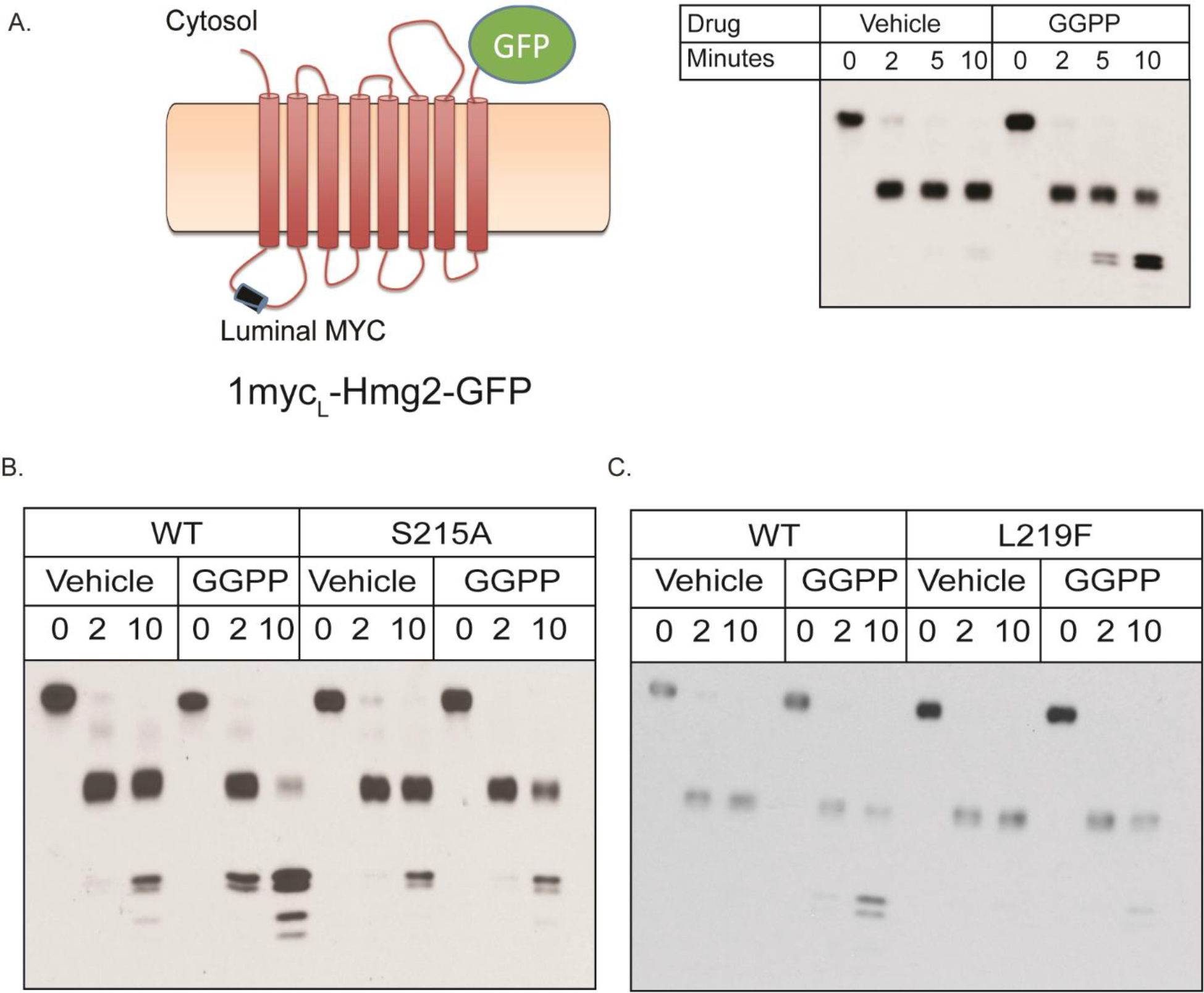
The SSD is required for GGPP-mediated mallosteric misfolding in vitro A. Left, schematic of the luminally tagged 1myc_L_-Hmg2-GFP reporter. A myc tag is inserted into the first luminal loop of Hmg2 and the Hmg2 transmembrane region is fused to GFP. (Cartoon first appeared in (18); used by permission) ER/Golgi microsomes were isolated from strains expressing 1myc_L_-Hmg2-GFP, treated with vehicle or GGPP, and subjected to proteolysis with trypsin. Addition of GGPP accelerates the rate of proteolysis approximately five-fold. B, C Stabilizing SSD mutations blocked GGPP-induced misfolding. Microsomes isolated from strains expressing wild-type or S215A 1myc_L_-Hmg2-GFP (B), or L219F 1myc_L_-Hmg2-GFP, were treated with vehicle or GGPP prior to proteolysis, and then treated with trypsin for the indicated times followed by immunoblotting for the luminal myc tag. GGPP induced misfolding and increased proteolysis in the wild-type protein, but was essentially ineffective

### INSIG independence of SSD-dependent mallostery

The SSD is perhaps best known for mediating the sterol-dependent binding to the regulatory proteins known as INSIGs (insulin-stimulated genes). In many cases, an SSD-INSIG interaction underlies regulation that requires an SSD (13, 29–33). For example, mammalian HMGcR requires SSD-mediated INSIG binding for sterol-regulated degradation (29, 30). Similarly, mammalian SCAP employs SSD-dependent INSIG binding to control the sterol regulated trafficking that mediates sterol-based control of transcription (33). S. cerevisiae has two INSIG homologs called Nsg1 and Nsg2. Nsg1 binds to Hmg2 in a sterol-dependent manner that strongly inhibits its regulated degradation (31, 32). Accordingly, we wondered if the SSD dependent, GGPP-stimulated degradation of Hmg2 was in any way dependent on either of the yeast INSIG homologs. Typically, we study Hmg2 regulated degradation at levels of Hmg2 in excess of native promoter Nsg expression (31). Nevertheless, it was formally possible that the Nsgs were involved in SSD-dependent degradation of Hmg2, so we directly tested that possibility.

Simultaneous deletion of Nsg1 and Nsg2 did not affect the steady state level of Hmg2-GFP as measured by flow cytometry (Fig. 3A, left). Furthermore, Hmg2-GFP remained normally responsive to GGPP in *nsg1Δnsg2Δ* double null strains (Fig. 3A, right). As a control, we also tested whether INSIG deletion affected the strong acquired stability of the SSD mutant S215A. As expected, S215A Hmg2-GFP levels were unaffected by deletion of both INSIGS and it remained stable and unresponsive to GGPP in the *nsg1Δnsg2Δ* strain (Fig. 3B, left and right, respectively). As expected from our earlier studies, expression of Nsg1 from the same strong promoter as that used for Hmg2-GFP, allowing stoichiometric interaction between the Nsg1 and Hmg2, increased the steady state level of Hmg2-GFP (Fig. 3C, left), and blocked the degradation-stimulating effects of GGPP (Fig. 3C, right). We next confirmed that INSIGs were not required for the SSD-dependent response to GGPP using the *in vitro* proteolysis assay. When 1myc_L_-Hmg2-GFP was expressed in the *nsg1Δnsg2Δ* double null strain, it responded normally to GGPP (Fig. 3D, left). Conversely, the presence of high levels of Nsg1 completely blocked GGPP-dependent misfolding of 1myc-Hmg2-GFP (Fig. 3D, right). Taken together, these data suggest that yeast INSIGs were not required for SSD-dependent regulated Hmg2 misfolding, but instead block the function of the SSD by binding, which we have shown occurs in a sterol-dependent manner (34). We have previously posited that this combination of regulatory effects allows for “contingency regulation” of Hmg2 ERAD, in which high GGPP and low sterols are the metabolic conditions required for Hmg2 degradation; appropriate for the natural physiology of HMGR in yeast (32). Whatever the evolutionary function of Nsg-mediated blockade of Hmg2 degradation, it is clear that GGPP-dependent mallostery of Hmg2 represents an autonomous, but INSIGs -modulated, physiological function of the Hmg2 SSD.

**Figure 3.**
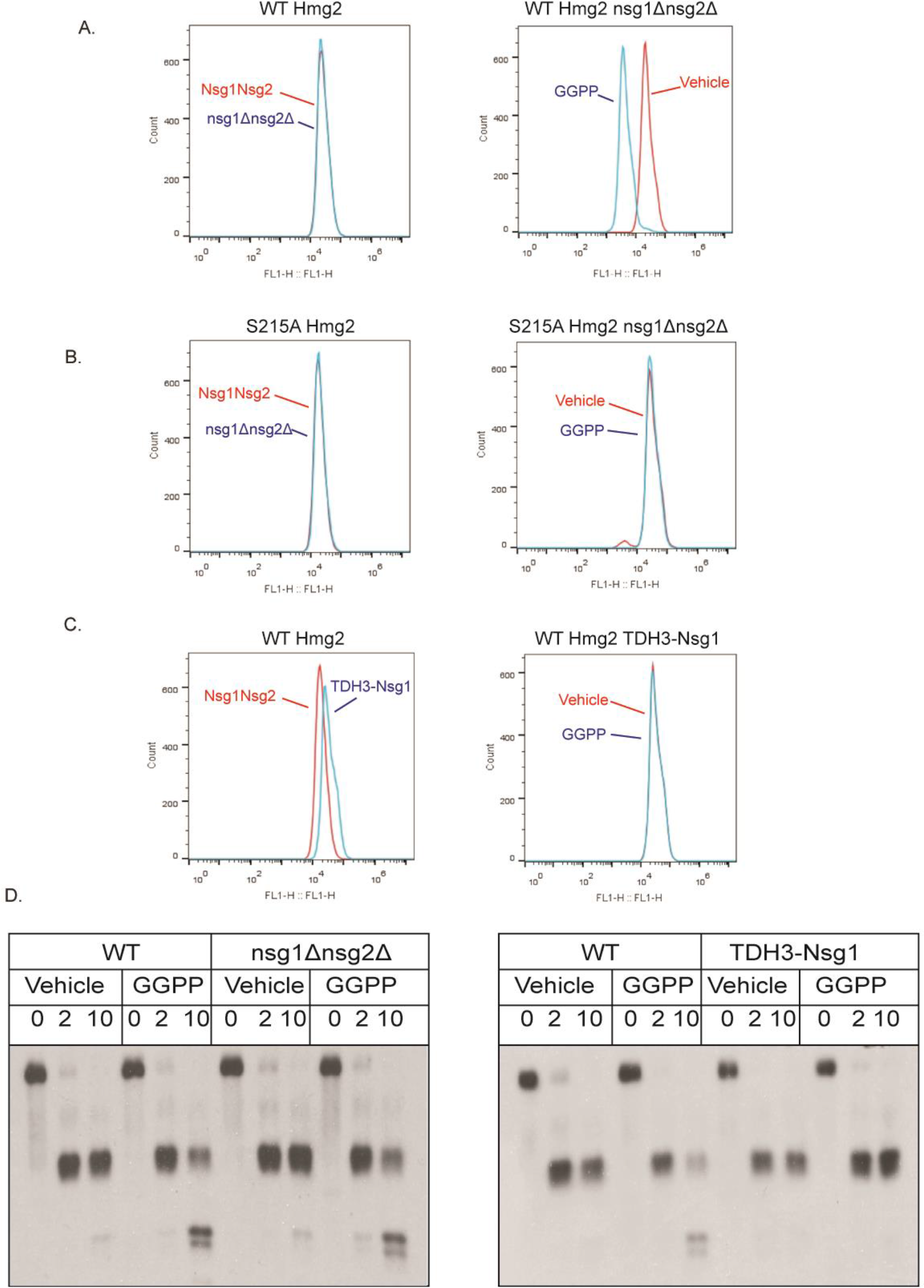
GGPP-regulated Hmg2 mallosteric misfolding did not require INSIG proteins, and was inhibited by excess Nsg1 A. Absence of yeast INSIGS Nsg1 and Nsg2 did not affect regulation degradation of Hmg2-GFP. Left panel, steady state levels of Hmg2-GFP in a strain with wild-type INSIG proteins (red curve) or an otherwise identical strain with missing both INSIG genes by virtue of nsg1Δnsg2Δ double null (blue curve). Right panel: in the INSIG double null, GGPP treatment caused expected drop in steady-state Hmg2-GFP levels (blue curve) compared to vehicle (red curve). B. Deletion of yeast INSIGs did not alter the stability or GGPP-non-responsiveness of the S215A mutant. Left panel, S215A Hmg2-GFP levels in a wild-type (red curve) or nsg1Δnsg2Δ double null strain (blue curve). Right panel, lack of GGPP responsiveness of S215A mutant in an nsg1Δnsg2Δ strain treated with vehicle (red) or GGPP (blue) for one hour. The block in regulation of the S215A mutant was not affected by INSIG deletion. C. Overexpression of Nsg1 increased Hmg2-GFP levels and prevented GGPP-induced degradation. Left, wild-type Hmg2-GFP expressed in a wild-type strain (red) or a strain producing Nsg1 from the strong TDH3 promoter (blue) were evaluated for steady-state fluorescence by flow cytometry. Right, cells expressing Hmg2-GFP in the an identical strain expressing Nsg1 from the strong TDH3 promoter were treated with vehicle (red) or GGPP (blue) and evaluated for steady state fluorescence by flow cytometry for one hour D. GGPP-induced in vitro misfolding of Hmg2-GFP did not require INSIGs, but was blocked by strong Nsg1 co-expression. Left, in vitro proteolysis of 1myc_L_-Hmg2-GFP expressed in wild-type and nsg1Δnsg2Δ yeast. GGPP induced misfolding and increased the rate of proteolysis in both backgrounds. Right, in vitro proteolysis of 1myc_L_-Hmg2-GFP expressed in a strain wild-type for INSIGs and a strain expression Nsg1 from the strong TDH3 promoter. Co-overexpression of Nsg1 blocked GGPP-induced misfolding.

### Separate determinants of Hmg2 mallosteric misfolding and ER degradation

We have previously identified a number of stabilizing mutations in Hmg2 (19, 35). Because regulated degradation of Hmg2 entails ligand-mediated misfolding followed by Hrd1-dependent ubiquitination and degradation, we wondered if any of the various mutants might lesion different parts of this multi-step regulatory process. Using the limited proteolysis assay to look specifically at the ligand-regulated misfolding step of Hmg2 regulation, we tested other stabilizing Hmg2 mutations to learn whether they, like conserved SSD residues, alter mallosteric misfolding, or other aspects of Hmg2 ERAD.

We focused on two lysines involved in Hmg2 degradation, K6 and K357, discovered in our early work (36). Each is required for Hmg2 regulated degradation: mutation of either to arginine results in complete stabilization at any concentration of the degradation signal (Fig. 4A) (17, 36, 37). Neither stabilized K → R mutant undergoes ubiquitination in vivo. As these lysines both face the cytosol, we had originally hypothesized that they might be ubiquitination sites, albethey quite distant along the primary sequence of Hmg2. Interestingly, human HMGcR similarly has two distant cytoplasmic lysines each required for regulated ERAD of the human form of the enzyme (22, 30).

**Figure 4.**
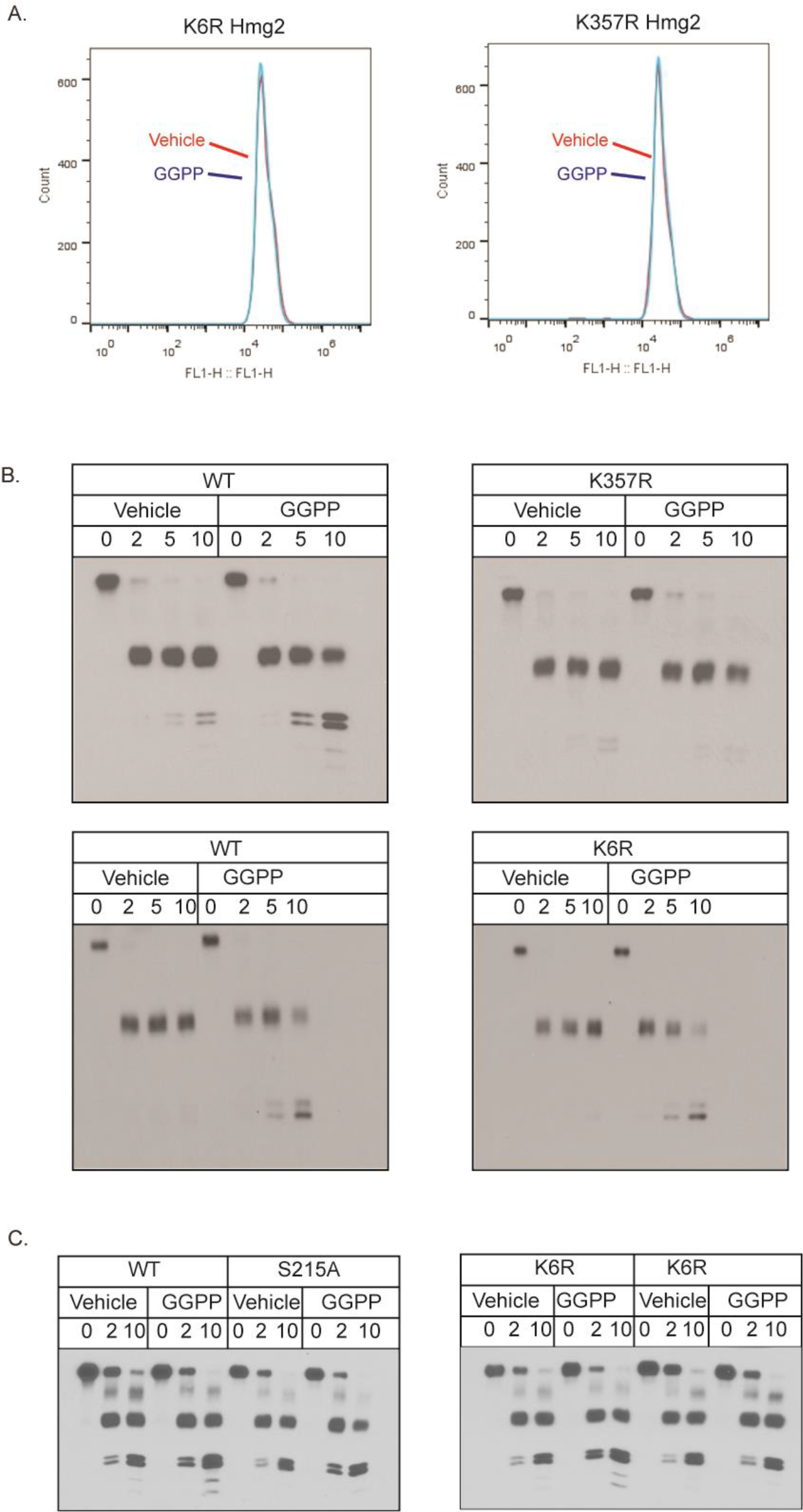
Separate sequence determinants of misfolding and degradation A. The two lysine-to-arginine mutations, K6R and K357R each rendered Hmg2-GFP insensitive to the degradation signal GGPP. Treatment strains expressing each mutant were treated with 22μM GGPP (blue curves) followed by 2 hrs incubation, and then subjected to flow cytometery. Unlike WT Hmg2-GFP (Fig 1C) did not t affect K6R or K357R Hmg2-GFP levels compared to vehicle treatment (red curves). B. The mutation K357R blocked GGPP-induced misfolding in vitro, but the K6R mutation did not. Microsomes expressing wild-type, K357R, and K6R 1myc_L_-Hmg2-GFP were isolated and treated with vehicle or 22μM GGPP prior to proteolysis as in earlier figures. GGPP did not cause misfolding and increased proteolysis in the K357R mutant (top panels). Conversely, K6R1myc_L_-Hmg2-GFP responded to GGPP like the wild-type protein (bottom panels). C. In-cis epistasis of the K6R mutation and the SSD mutation S215A. Both wild-type and K6R 1myc_L_-Hmg2-GFP misfold upon 22 μM GGPP treatment, while the K6R S215A double mutation did not respond to GGPP treatment.

We revisited these stabilizing mutations and tested each with the in vitro proteolysis assay to evaluate their ability to support GGPP-regulated misfolding. Despite not being a conserved SSD residue, K357R 1myc_L_-Hmg2-GFP behaved like the SSD mutants discussed above: it was completely unresponsive to added GGPP (Fig. 4B). Conversely, K6R 1myc_L_-Hmg2-GFP, despite its extreme in vivo stability, showed an entirely normal response to GGPP, behaving identically to wild type 1myc_L_-Hmg2-GFP in the misfolding assay (Fig. 4B). This defined for the first time two classes of determinant for Hmg2 degradation, and an apparent sequence of events: GGPP dependent misfolding that was affected by SSD mutants such as S215A, or K357R, followed by Hrd1-dependent ubiquitination that required K6, but after SSD-dependent misfolding. To test this idea, we performed a “cis-epigenetic” test of this model with double mutants from each class: the K6R mutant that allows regulated misfolding, and our strongest, best characterized SSD mutant—S215A—that fails to undergo regulated misfolding. If misfolding precedes ubiquitination, we would expect the K6R/S215A double mutant to lose GGPP dependent misfolding and remain stable, that is, the S215A would be expected to be epistatic to the K6R mutant, which is what we observed. (Fig. 4C).

### SSD functions in the kinetics of GGPP-dependent Hmg2 misfolding

The above demonstrations of a separable, independent role of the SSD led us to more fully investigate the nature of the SSDs action in Hmg2 mallosteric misfolding. We wondered if the SSD was involved in determining the rate or extent of response to GGPP. Specifically, we examined whether Hmg2 with a strong stabilizing SSD mutation could still respond to GGPP at sufficiently high concentrations or long incubation times. We performed time course experiments testing wild-type or S215A Hmg2-GFP in the in vitro proteolysis assay at a variety of GGPP concentrations and incubation times. Although the stable SSD mutants had not responded to GGPP in any of our normally conducted assays, we found that when incubated overnight with a high concentration of GGPP (approximately 1000 times that required for initial effects in the wild type protein), S215A 1myc-Hmg2-GFP did indeed respond to GGPP (Fig. 5A). Despite the apparent much lower potency, the effect remained highly specific for GGPP: the previously described inactive analog 2F-GGPP was similarly unable to affect S215A Hmg2 in the extended time assay (Fig. 5B). A time course of GGPP incubation confirmed that the response time of S215A 1myc_L_-Hmg2-GFP was severely delayed, starting to respond to GGPP only after 2.5 hours of treatment with a maximal response occurring by 3.5 hours (Fig. 5C). For comparison, wild type 1myc-Hmg2-GFP responds to signal almost immediately; within <5 minutes when GGPP is added at the same time as protease.

**Figure 5.**
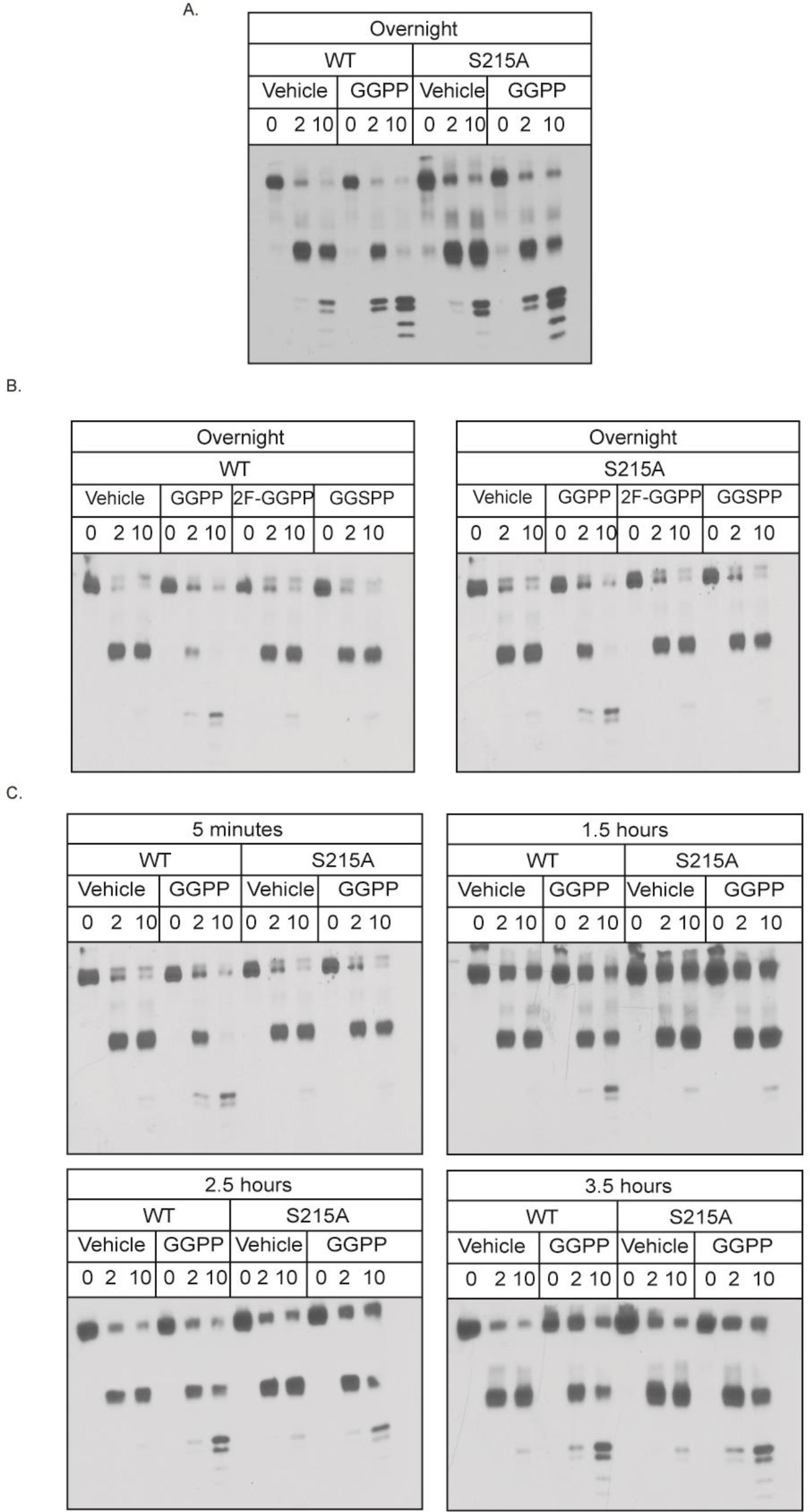
GGPP caused mallosteric misfolding of the stable SSD mutant S215A at high concentrations when treated for long time courses A. Overnight GGPP treatment caused in vitro misfolding of S215A Hmg2. Western blots of in vitro proteolysis performed on membranes from cells expressing wild-type or S215A 1myc_L_-Hmg2-GFP and incubated with vehicle or 22 μM GGPP overnight (15 hours). B. GGPP-mediated, long time course misfolding of S215A misfolding maintained high GGPP structural specificity observed in WT Hmg2. Specificity of overnight misfolding of wild-type or S215A 1myc_L_-Hmg2-GFP was tested by treatment with 22 μM of the close analogs of GGPP, 2-fluouro GGPP (2F-GGPP) and s-thiolo GGPP (GGSPP), shown to be inactive in more canonical rapid structural changes in wt Hmg2. Neither caused misfolding of either wild-type or S215A Hmg2 at the high concentrations and long time courses employed in this experiment C. Time course of wild-type and S215A GGPP-induced misfolding. Wild-type 1myc_L_-Hmg2-GFP misfolded in response to GGPP more quickly than can be measured, within 5 minutes. S215A 1myc_L_-Hmg2-GFP began to misfold in response to GGPP after approximately 2.5 hours of treatment.

### “Toxic subunit” effects as a test of the Hmg2 multimer in mallostery

We had previously shown that the GGPP-mediated reversible misfolding of Hmg2 has many attributes of allosteric regulation (18). These include: high ligand potency and specificity, the existence of a specific GGPP antagonist, reversibility, and the biochemical observation that the Hmg2 transmembrane domain is a multimer in vivo. These similarities led us to suggest the name “mallostery” to describe GGPP-mediated misfolding of Hmg2. Nearly all cases of allosteric regulation involve structural changes based on concerted interaction between subunits of a regulated multimer. Accordingly, we wondered if the GGPP-mediated mallosteric misfolding would similarly depend on the observed multimeric structure of the Hmg2 transmembrane domain. The existence of the above-described mutants deficient in GGPP-mediated misfolding provided a powerful new experimental tool to test this feature of the mallosteric model. The idea is based on the observation of dominant negative mutations in many allosteric and cooperative proteins, in which the presence of a non-responsive mutant subunit within an allosteric multimer can block the regulation of the whole mulitimeric assembly ((38, 39). We hypothesized that the presence of a non-responsive SSD mutant Hmg2 present as a subunit of an Hmg2 multimer would interfere with GGPP-mediated response of the mixed quaternary structure. Conversely, we predicted that the K6R mutant, that still undergoes GGPP stimulated misfolding but is not ubiquitinated, would still permit regulated degradation of its wild-type partners within a mixed multimer. To test this idea, we constructed yeast strains co-expressing both wild-type and non-responsive SSD mutants, such that the behavior of the wild-type protein could be independently examined in the presence of co-expressed mutant or wild-type controls. Specifically, we co-expressed wild-type, optically detectable, normally regulated Hmg2-GFP, along with non-fluorescent, non-responsive S215A as well as control strains with a wild-type version of the non-fluorescent co-expressee. In this way we examined the behavior of only the normally regulated Hmg2-GFP by optical means when in the presence or absence of the co-expressed, non-fluorescent, non-responsive mutants (Fig. 6A).

**Figure 6.**
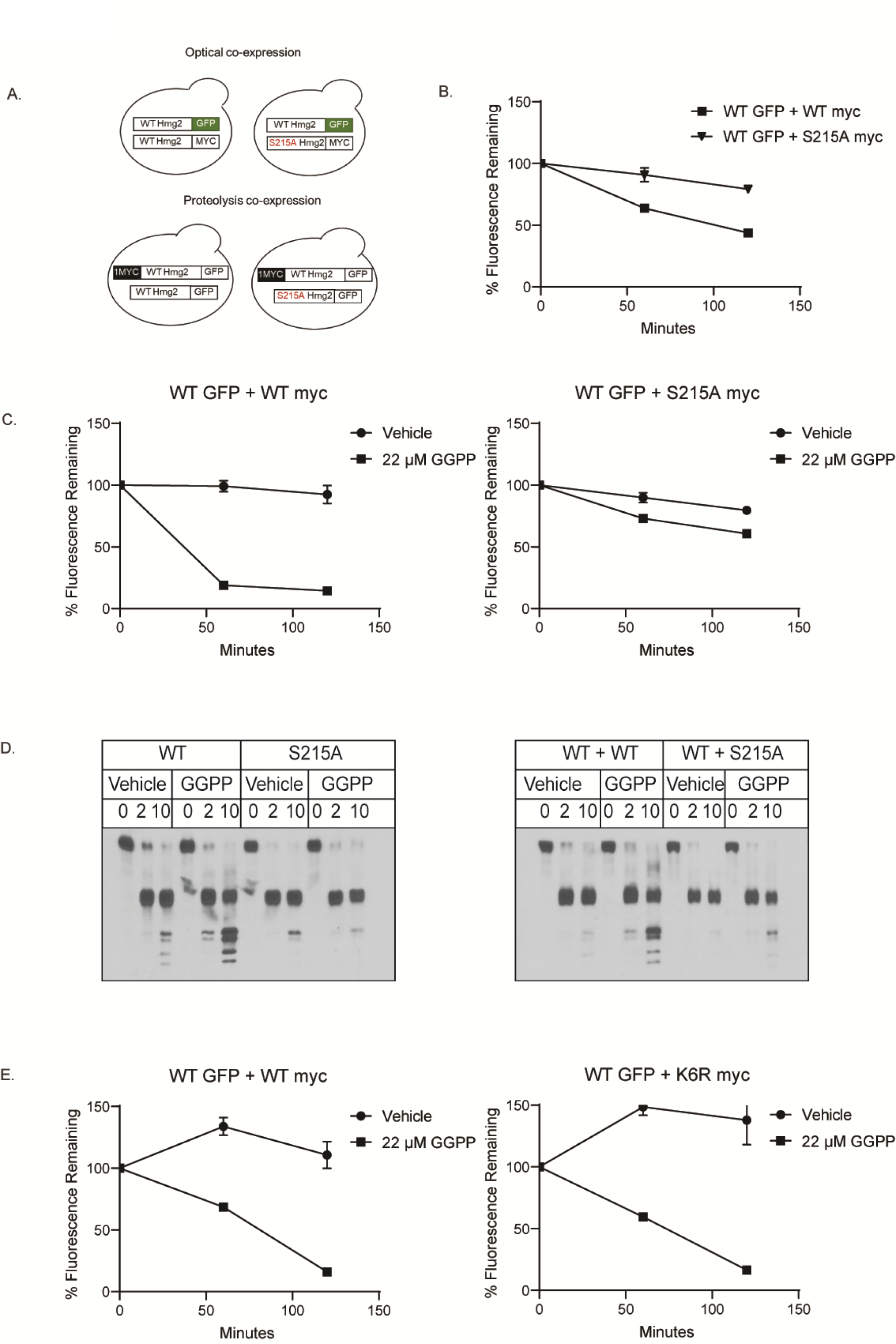
Trans effects of mutated SSDs on regulated misfolding A. Cartoon showing experimental setup for co-expression experiments. For optical experiments, WT Hmg2-GFP was co-expressed alongside a non-fluorescent myc-tagged mutant S215A or WT (control) copy (top). For proteolysis experiments, WT 1myc_L_-Hmg2-GFP was co-expressed alongside a GFP-tagged, without myc, mutant S215A 1myc_L_-Hmg2-GFP or WT (control) (bottom). B. Co-expression of dark (non-fluoresecent), myc-tagged, non-mallosteric mutant S215A Hmg2 strongly stabilized wild-type Hmg2-GFP. Wild-type Hmg2-GFP was expressed along with dark wild-type Hmg2-myc (filled squares) or S215A Hmg2-myc (filled triangles). In a cycloheximide chase, the dark S215A Hmg2-myc slowed the degradation of wild-type Hmg2-GFP when compared to a strain co-expressing wild-type Hmg2-myc. Error bars are SEM. C. Co-expression of the dark S215A Hmg2-myc inhibited the response of wild-type Hmg2-GFP to GGPP. Wild-type Hmg2-GFP was co-expressed with wild-type Hmg2-myc (left) or S215A Hmg2-myc (right). Co-expression of the mutated Hmg2-myc which cannot misfold, but not the wild-type Hmg2-myc, partially blocked GGPP-induced degradation of the wild-type Hmg2-GFP. Filled circles show vehicle control, and filled squares show GGPP treatment. Error bars are SEM. D. Co-expression of a distinct non-mallosteric SSD mutant Hmg2-GFP strongly blocked GGPP-induced misfolding of wild-type 1myc_L_-Hmg2-GFP in vitro. Left, GGPP induced misfolding and caused increased proteolytic cleavage in wild-type but not S215A 1myc_L_-Hmg2-GFP. Right, co-expression of wild-type Hmg2-GFP without a a myc tag did not interfere proteolysis of wild-type 1myc_L_-Hmg2-GFP. However, co-expression of S215A Hmg2-GFP with no myc tag attenuated GGPP-induced misfolding of the wild-type 1myc_L_-Hmg2-GFP copy. E. Co-expression of the highly stable but still mallosteric K6R Hmg2-myc, allowed normal Hmg2-GFP regulation. Cells expressing Hmg2-GFP and co-expressing a dark wild-type Hmg2-myc (left) or K6R Hmg2-myc (right) were treated with vehicle (filled circles) or GGPP (filled squares), and Hmg2-GFP levels were assayed by flow cytometry. The K6R Hmg2-myc did not block regulation of the wild-type Hmg2-GFP. Error bars are SEM.

When these two versions of Hmg2 were co-expressed, the wild type Hmg2-GFP co-expressed with K357R was degraded more slowly in a cycloheximide chase, implying that the co-expressed dark, K357R mutant stabilized the fluorescent wild type Hmg2-GFP (Fig. 6B). Similarly, addition of GGPP to the strain co-expressing non-responsive K357R-Hmg2 was much less effective at causing degradation of the wild-type Hmg2-GFP than in the otherwise identical strain co-expressing wild type Hmg2 (Fig. 6C).

We also tested for intersubunit interactions using the in vitro limited proteolysis assay. For these experiments, we used wild type 1myc_L_-Hmg2-GFP from the limited proteolysis experiments above, co-expressed with either wild-type or S215A Hmg2-GFP with no lumenal myc tag, so that, in this assay as well, we could selectively observe the proteolytic response of only 1myc_L_-Hmg2-GFP in the presence of co-expressed but not-myc-tagged Hmg2 mutants (6D). We included a strain expressing the single, highly stable protein S215A-1myc_L_-Hmg2-GFP as a positive control to evaluate the strength of any trans effects observed. Again, we found that co-expression of the non-responsive S215A Hmg2-GFP interfered with the ability of the wild type copy to undergo GGPP-dependent misfolding (6D, right panel). Wild type 1myc_L_-Hmg2-GFP underwent normal in vitro misfolding in response to GGPP when the co-expressed, non-myc tagged test protein was wild type, but its response to GGPP was severely blunted when the co-expressed, non-tagged protein had the S215A mutation (Fig. 6D). In fact, the in-trans effect of a non-fluorescent S215A mutant on co-expressed wt Hmg2-GFP was as strong in this assay as the effect of the mutation on the lone expressed S215A-Hmg2-GFP (compare 6D right side pairs of left (cis S215A effect) and right (co-expressed S215A effect) panels). Taken together, these in vivo and in vitro studies indicated that the mallosteric action of the SSD operates at the level of multimeric Hmg2, as indicated by the ability of non-responsive SSD mutants to strongly affect the response of co-expressed but normal Hmg2 to the GGPP signal in vivo or in vitro.

The above co-expression tests of allosteric-style communication between subunits required for a “toxic subunit” effect is both powerful and classic, being borrowed from the deeply rooted literature of allosteric regulation (38, 39). Our model is that the regulated misfolding caused by an intact, wild-type SSD operates through the quaternary structure of the Hmg2 transmembrane region, and the presence of non-responsive SSDs drastically affects the GGPP-mediated changes in structure of the multimer. However, other models could explain the interference of normal degradation caused by a co-expressed, highly stable version of Hmg2. Accordingly, we ran a critical control co-expression test that further capitalized on the mutant analysis above. We tested if the still-GGPP-responsive but highly stable K6R mutant had any effect when co-expressed with normal regulated Hmg2-GFP. The K6R mutant is also strongly stabilized like the S215A or K357R mutants, but underwent normal GGPP-mediated structural changes in vitro as shown above. Unlike the SSD mutants, coexpressing the highly stable K6R mutant had no effect on the degradation or the GGPP response of also-present Hmg2-GFP, indicating that the effect of the similarly stable but non-functional SSD mutants was specifically due to the loss of regulated misfolding within the co-expressed multimers (Fig. 6D).

Taken together, these studies strongly imply that the “mallosteric model” of GGPP causing reversible misfolding through concerted structural changes in the quaternary Hmg2 transmembrane domain is the mechanism of ligand regulated Hmg2 misfolding.

### GGPP-dependent mallostery was membrane-autonomous

Finally, we tested the ability of the Hmg2 to undergo GGPP-dependent misfolding when removed from its normal membrane context, after detergent solublization. We prepared microsomes from the *in vitro* proteolysis strain expressing 1myc_L_-Hmg2-GFP and subjected them to solubilization with a variety of detergents. Three of the detergents tested, Fos-Choline-13, Decyl Maltose Neopentyl Glycol (DMNG), and digitonin yielded soluble 1myc_L_-Hmg2-GFP that retained its normal pattern of proteolytic cleavage (Fig. 7A). This solubilized Hmg2 was, however, more sensitive to trypsin (data not shown), so lower concentrations of the protease were used in these experiments.

**Figure 7.**
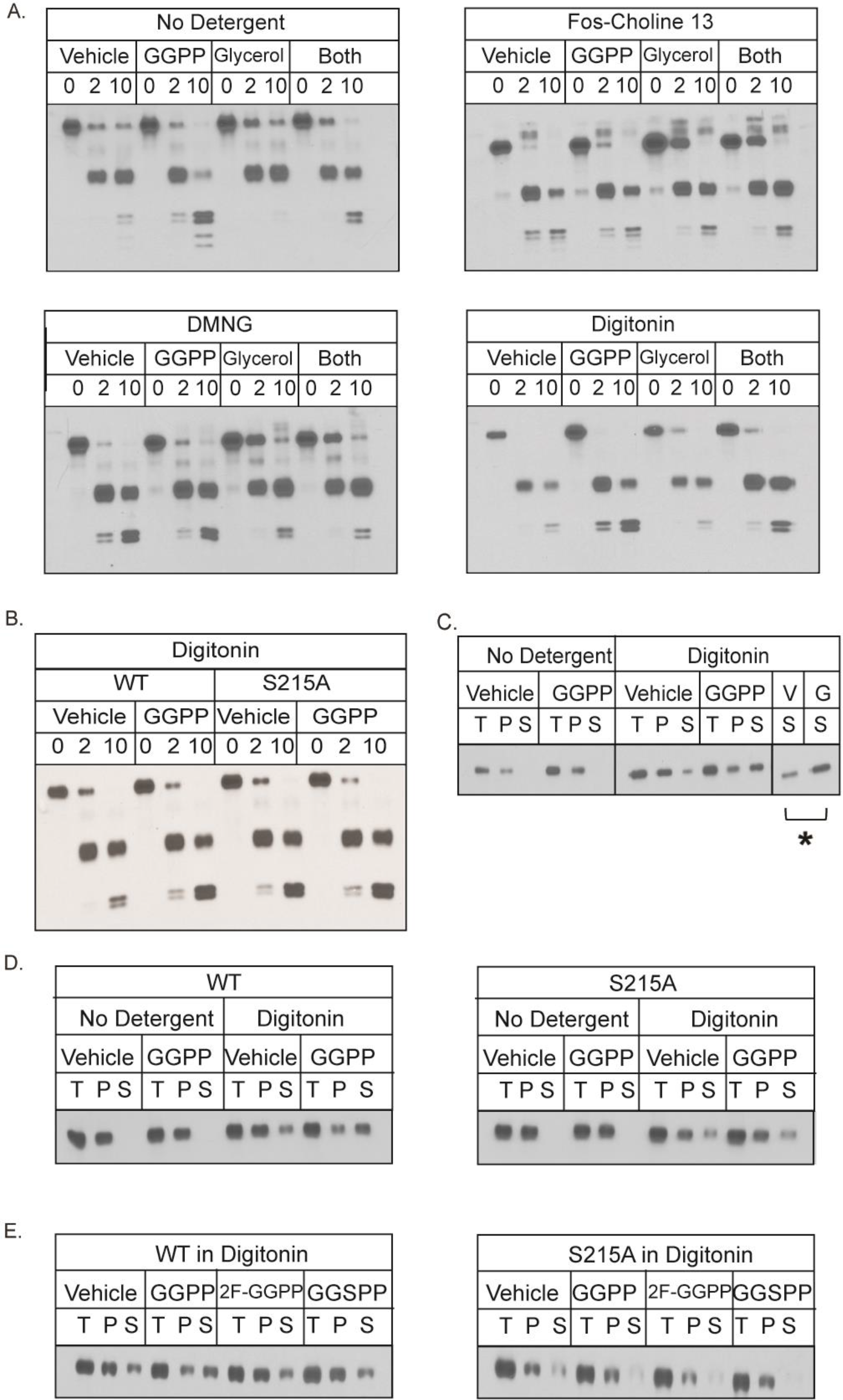
In vitro proteolysis assay of micellar 1myc_L_-Hmg2-GFP is intact, and responses to glycerol and GGPP A. The non-ionic detergents fos-choline 13, decyl maltose neopentyl glycol (DMNG), and digitonin allowed the time dependent proteolytic cleavage pattern of 1myc_L_-Hmg2-GFP in solution. Microsomes from cells expressing 1myc_L_-Hmg2-GFP were isolated as previously and left unsolubilized (top left) or subjected to solubilization with fos-choline 13 (top right), DMNG (bottom left), or digitonin (bottom right) as described in text. For the no detergent condition, the microsome pellet was used; for the three detergent conditions, solubilized microsomes were clarified by ultracentrifugation and the supernatant was subjected to proteolysis. In fos-choline 13, DMNG, and digitonin, the 1myc_L_-Hmg2-GFP myc tag remained intact during proteolysis. 1myc_L_-Hmg2-GFP remained responsive to the action of the chemical chaperon glycerol in all three detergents as well. Furthermore, 1myc_L_-Hmg2-GFP solubilized in digitonin remained responsive to GGPP in vitro B. Digitonin solubilization preserved the SSD requirement for 1myc_L_-Hmg2-GFP misfolding. When solubilized with digitonin, proteolysis of wild-type 1myc_L_-Hmg2-GFP increased in response to GGPP treatment, but proteolysis of S215A 1myc_L_-Hmg2-GFP did not. C. GGPP treatment during solubilization increased the solubility of 1myc_L_-Hmg2-GFP. Left, when not solubilized and subjected to centrifugation, 1myc_L_-Hmg2-GFP was present only in the pellet when detected by western blotting for the myc tag. GGPP did not affect 1myc_L_-Hmg2- GFP fractionation in unsolubilized microsomes. Right, when microsomes were solubilized with digitonin, 1myc_L_-Hmg2-GFP was present in both pellet and supernatant fractions. Treatment of microsomes with 22 μM GGPP during solubilization increased the amount of 1myc_L_-Hmg2-GFP in the supernatant fraction. Far right, comparison of the amount of 1myc_L_-Hmg2-GFP in the supernatant when treated with vehicle versus GGPP. *denotes p ≤ 0.05. D. The solubilization effect of added GGPP was SSD dependent. Left, treating preparations with GGPP during solubilization increased the amount of wild-type 1myc_L_-Hmg2-GFP detectable in the supernatant. However, the solubility of S215A 1myc_L_-Hmg2-GFP was not affected by GGPP. E. The increase in solubilization was specific for GGPP. Treating preparations with the close analogues of GGPP, 2-fluoro-GGPP (2F-GGPP) or s-thiolo GGPP (GGSPP) did not increase the amount of 1myc_L_-Hmg2-GFP detectable in the supernatant.

The preservation of the pattern of proteolysis was somewhat surprising, given that we had assumed the luminal location of the 1myc_L_ tag was required for protection from the added trypsin. These results imply that the luminal tag is further protected by some aspect of Hmg2 structure independent of lumenal isolation. Whatever the cause of this “solution autonomy”, the preservation of the myc-detected proteolysis pattern allowed us to examine the role of the mallosteric behavior of Hmg2 when completely separate from the ER membrane. Accordingly, we tested the response of detergent solubilized 1myc_L_-Hmg2-GFP to either chemical chaperones or GGPP using a soluble variant of the microsomal limited proteolysis assay. Solubilized 1myc_L_-Hmg2-GFP appeared to be constitutively more structurally open than when present in the ER membrane, with high basal rates of proteolysis. However, it retained the ability to respond to the chemical chaperone glycerol in all three detergents: preparations treated with 20% glycerol were more resistant to proteolysis, as in the normal microsomal assay (Fig. 7A va B,C,D). The GGPP dependent structural response was more sensitive to choice of solubilization detergent. Hmg2-GFP solubilized with either Fos-Choline-13 or DMNG had no detectable response to GGPP, whether alone or in combination with glycerol treatment (Fig. 7A). Conversely, digitonin-solubilized preparations did indeed respond to GGPP in the limited proteolysis assay, although at somewhat higher concentrations than required in the original microsomal assay (Fig. 7A). Importantly, the nonresponding stable S215A Hmg2 did not respond to GGPP in the digitonin-solubilized state (Fig. 7B), indicating that the SSD mediates the GGPP response in the micellar state in the absence of any membrane. Furthermore, the degree of response to GGPP in the digitonin solubilized Hmg2 was lessened by co-incubation with glycerol, as is the case in the microsomal assay as well as in vivo.

In the course of these studies, we also found that GGPP affected Hmg2 detergent solubility. After preparing and solubilizing microsomes, we added vehicle or GGPP to the preparations and further incubated for 1 hour with gentle shaking. Afterward, we separated the samples by ultracentrifugation, and found that preparations treated with GGPP had been more effectively solubilized by digitonin (Fig. 7C). The stable S215A Hmg2 did not become more solubilized by GGPP in these experiments, indicating that this effect is specific both to the structure of GGPP and to the misfolding capability of the SSD (Fig. 7D). Furthermore, the inactive analog of GGPP, 2F-GGPP, which does not stimulate Hmg2 degradation in vivo or misfolding in vitro, did not have any effect on solubility in this assay (Fig. 7E). Although these effects are small, it is intriguing that a straightforward, reproducible, and highly specific readout of the mallosteric response manaifests at the level of protein biochemical behavior. Importantly, the change in ratio of soluble Hmg2 (S) to total Hmg2 (T) was statistically significant, as indicated by the bar in Fig 7C

The whole of the above data indicate that the SSD is required for regulated misfolding and degradation of Hmg2. Lesions in the SSD block Hmg2 from responding to the degradation signal GGPP, preventing in vivo degradation and in vitro misfolding. The role of the SSD appears to be “kinetic,” with long incubations at extremely high concentrations of signal eventually overcoming its loss. Hmg2 forms multimeric structures, and mutations in the SSD can block regulation in wild type Hmg2 *in-trans*. The SSD constitutes a separate determinant for Hmg2 regulated misfolding, and its function is autonomous of INSIG proteins and partially retained even when removed from the normal context of the ER membrane.

## Discussion

In this work we set out to understand structural features of Hmg2 that contribute to regulated degradation, with particular emphasis on role of the conserved sterol sensing domain (SSD) in the ligand-regulated misfolding that underlies mallostery. The SSD is a multispanning membrane motif found in a number of eukaryotic proteins involved in sterol synthesis, transport, and regulation. The demonstrated importance of the Hmg2 SSD in GGPP-mediated misfolding was consistent with a broad role of SSD in mediating protein structural changes as a mechanism of metabolic regulation (13, 16, 40). Importantly, our results demonstrate an autonomous role for a bone fide SSD in protein regulation independent of the frequently described INSIG-SSD interactions often required for SSD actions. Furthermore, the genetic separability of regulated misfolding and HRD-dependent ubiquitination that capitalized on conserved SSD residues allowed us to test the mallosteric model of GGPP-regulated misfolding using co-expression studies. Taken together, these data indicate that ligand-mediated, regulatory misfolding can show precise and evolvable sequence underpinnings that bode well for both understanding the function of the SSD, as well as opening the door for development of degradation-based therapeutic molecules for a variety of desired protein targets.

We tested highly conserved SSD residues for their role in regulated misfolding using both flow cytometry in vivo and a limited proteolysis assay of Hmg2 structure in vitro. Mutations in the SSD that stabilize Hmg2 in vivo also blocked GGPP-induced misfolding of Hmg2 in vitro, meaning that the proteolysis assay can serve as a biochemical test of SSD function. We had shown previously that all phenotypic SSD point mutations stabilize Hmg2 in vivo--no conserved residue mutants resulted in decreased stability--suggesting that the normal function of SSD is to allow regulated misfolding to occur. Because of the near-universal involvement of INSIGs in the action of SSDs in cholesterol regulation (13, 29, 31–33), we tested whether the SSD-dependent structural change was intrinsic to the domain itself, or rather required the yeast INSIG orthologs Nsg1 or Nsg2. Despite the expectation for an INSIG role, we found that neither INSIG paralog was required for SSD-dependent GGPP-regulated misfolding. However, consistent with our earlier studies (31, 32), coexpressing Nsg1 to levels matching Hmg2 actually blocked regulated in vitro misfolding and in vivo degradation. Hmg2 has been previously found to bind to INSIGs, but their role in yeast is an inhibitory one: yeast INSIGs blocks the ability of GGPP to promote Hmg2 degradation, and this interaction between Hmg2 and Nsg1 is lanosterol dependent. The model put forth from those studies is that Hmg2 undergoes GGPP regulated degradation, and that INSIGs allow this only to occur when sterol synthesis (and thus lanosterol levels) are low and isoprenes such as GGPP would tend to accumulate (34). The data above are consistent with those previous studies, since in vivo regulation of Hmg2 by GGPP occurs in the absence of INSIG proteins, and the SSD-dependent regulation of Hmg2 is the core molecular feature of this regulation. This INSIG-independence of SSD action establishes a clear, autonomous role for this conserved domain, which, in the case of Hmg2, is further regulated by INSIGs to impart a second layer of sterol pathway control.

It is worth noting that yeast Hmg2 occupies a particularly advantageous position for the study of SSDs and INSIGs. In most reported cases, either SSD action is totally dependent on INSIG proteins (SCAP; HMGcR regulation) (33, 41–43), or completely independent of INSIGs (NPC1, S. pombe SCAP) (44, 45). Yeast Hmg2 regulation occupies a mechanistic “sweet spot”, in which the SSD clearly functions autonomously to allow physiologically useful regulation by GGPP, but is modulated by sterol-dependent interaction with the endogenous INSIGs. Thus, questions of the independent and intertwined roles of these two key components of sterol biology, and how this interaction evolved, can be studied using the combined powers of yeast genetics, biochemistry and molecular biology.

Using our limited proteolysis assay of GGPP action, we tested other, non-SSD Hmg2-stabilizing mutations. Our early work had identified mutations in a pair of cytoplasmic lysines, K6, located on the cytoplasmic N-terminus of Hmg2, and K357, located in its sixth cytoplasmic loop. Both K6R Hmg2 and K357R Hmg2 are extremely stable in vivo, both in a cycloheximide chase or upon treatment of cells with the misfolding signal GGPP (35). In addition, unlike wild-type Hmg2, neither mutated protein is detectably ubiquitinated in response to GGPP treatment. Accordingly, we tested these mutations in our in vitro misfolding assay. We found that while K357R Hmg2 did not respond to GGPP, K6R Hmg2 still underwent normal GGPP-dependent misfolding in vitro, in a manner identical to the wild-type protein. This was the first time we have identified separable Hmg2 sequence determinants required for misfolding versus ubiquitination and degradation. Additionally, this result suggests a model wherein K6, as a cytosol-facing lysine which prevents Hmg2 ubiquitination and degradation but not misfolding, may be a major, or first ubiquitination site, while lysine 357 is instead (or in addition) required for regulated GGPP-dependent misfolding in response to GGPP. Indeed, studies with double mutants demonstrated that still-misfolded yet stable K6R mutant showed the expected SSD dependency in its response to GGPP, indicating the autonomy of these two aspects of GGPP-regulated degradation.

The evidence for an autonomous role of the SSD and the separability of that role from ubiquitination and degradation led us to further explore the action of the SSD in GGPP-regulated misfolding. We found through extended time course experiments that the highly conserved SSD residue mutant S215A-Hmg2, which is extremely stable in vivo and in vitro, could indeed responded to GGPP, but only after long incubation times, with a response substantially delayed compared to the response of the wild type protein. Remarkably, the delayed response of the S215A mutant still showed the high structural specificity of the GGPP stimulus: the inactive GGPP analogues remained inactive in the long-time course response of these stable variants. This indicates that the GGPP binding site may be distinct from a structure defined by the SSD, since the preservation of the strong structural specificity implies the still-unknown GGPP binding site is intact. Clearly, the next step will have to be a combination of binding and structural studies to resolve this intriguing question.

We further explored the role of Hmg2’s transmembrane region in the mallosteric response to the GGPP ligand by testing for trans-effects in co-expression experiments. In light of the demonstrable role for SSD in the effect of GGPP, and the fact that the Hmg2 transmembrane region exists as a multimer, we asked if the presence of subunits with non-responsive SSD mutations could affect the regulation of a wild-type Hmg2 within the same multimer through interactions typical of allostery. Specifically, we co-expressed mutant and wild-type SSDs in the same Hmg2 cells and measured the effect of the lesioned domain on the co-expressed wild-type domain’s ability to undergo GGPP-dependent misfolding and degradation. Indeed, we found that a non-responsive SSD is able to interfere with the action of the wild-type protein in trans. Importantly, a similarly stable, equally co-expressed K6R mutant, but which has a normal SSD-dependent response to GGPP, did not interfere with wild-type regulation. This sort of “poisoned subunit” experiment provides further evidence for SSD-mediated regulation of the Hmg2 folding state based on the mallosteric model. These co-expression experiments suggest a model for Hmg2 misfolding wherein the transmembrane domains of multimeric Hmg2 must cooperate to undergo concerted, multi-subunit based GGPP-regulated structural changes, which might be expected for the development of mallosteric regulation as a variation of cooperativity and allostery.

The above findings suggest a model for the SSD as an autonomous module for promoting misfolding in response to a regulatory ligand. By this model, GGPP binding at an unknown Hmg2 site causes the SSD to undergo a conformational change that renders the protein more susceptible to the HRD quality control pathway. Lesions in the SSD block this change at physiologically relevant concentrations and time scales. This role for the SSD appears to be a kinetic one, as mutations in the SSD do not appear to block the conformational change absolutely, nor the high specificity of the GGPP ligand, so much as render it impractically slow for cellular regulation.

The mallosteric model for SSD action leaves several open questions, first and foremost, what is the mechanistic role of the SSD? Is the SSD executing a structural transition between a folded and “misfolded” state in response to the misfolding signal GGPP, or alternatively, is the SSD responsible for GGPP binding? Our current in vitro approaches do not definitively distinguish between mutations in Hmg2 which eliminate GGPP binding and those which allow binding but disable misfolding in response to binding elsewhere. Our time course experiments showing that a mutated SSD can still specifically respond to GGPP, albeit over the course of several hours rather than seconds, suggest that the SSD may play a role in allowing rapid transition between a folded and misfolded state, rather than mediating binding per se. The idea that SSD mediates structural changes in response to binding at a distinct site has arisen in studies of SCAP, which responds to sterols that bind to the first luminal loop of the protein by an SSD-dependent structural change (46, 47). Intriguingly, sequence and structural homology to SCAP and small regions of GGPP binding proteins suggest a similar distinct GGPP binding site, although direct tests have not yet been performed.

Alternatively, there are also studies indicating SSDs can directly interact with ligands, and thus allow the possibility that it is the GGPP binding site (48–52). The SSD is related to the Resistance Nodulation Division (RND) domain, which is conserved in all domains of life and found in many transporters (53–56). Indeed, recent structures of the SSD of NPC1 suggest that in that protein the SSD functions in transport of sterol-related molecules, and have located a putative sterol-interacting pocket within the domain (51), lending credence to the alternate possibility of the SSD as the GGPP binding site. Clearly further studies are required to resolve the divergent mechanistic underpinning of the mallosteric regulation of Hmg2 ERAD.

In the course of this work, we extended our analysis of GGPP-mediated misfolding to detergent-solubilized Hmg2. We wondered if solubilized 1myc_L_-Hmg2-GFP would still show a useful proteolytic patter in the absence of a protected luminal space, and given that, if detergent solubilized protein would be remain responsive to chemical chaperones and/or GGPP. Remarkably, we found that all three of these responses could be true for Hmg2 in solution: the overall proteolytic pattern of Hmg2 in the in vitro misfolding assay was preserved when solubilized in several weak detergents, and Hmg2 in detergent solution retained the ability to be chaperoned by glycerol. Furthermore, in digitonin Hmg2 retained its ability to undergo GGPP-mediated misfolding, with a preserved requirement for an intact SSD, and the same high structural dependence on the GGPP molecule. The ability of solubilized Hmg2 to undergo ligand-regulated structural changes heralds a new collection of approaches that could resolve nearly all of the structural and functional questions put forth above about the role of SSS in mallosteric regulation of Hmg2, and the molecular functions of SSD.

The SSD is conserved in a variety of proteins from yeast to humans (13, 23, 56–61), many of them implicated in human diseases, ranging from dyslipidemia, which affects nearly half of American adults and a quarter of American youths in some fashion (62, 63), to inborn genetic disorders (64, 65), to cancer (66). The human homolog of Hmg2, HMGR, as well as the SSD-containing SCAP, are both critical components of sterol biosynthesis and regulation, and human HMGR is the target of cholesterol-lowering statin class of drugs. Another SSD-containing protein, NPC1-Like Protein 1 (NPC1L1) is involved in enteric cholesterol transport and targeted by the cholesterol-lowing drug ezetimibe (67, 68). Furthermore, three proteins associated with human inborn diseases, 7-dehydrocholesterol reductase (DHCR7), lesions in which cause Smith-Lemli-Opitz Syndrome (69–71), NPC1, lesions in which cause Niemann Pick Disease Type C (72, 73), and Patched, a regulator of Hedgehog signaling associated with the human cancer disorder Gorlin Syndrome (66, 74), all contain SSDs. In many proteins where the role of the SSD is well-understood, the motif seems to allow molecular outcomes related to protein quality-control, ranging from entry into the ERAD pathway, as in both *S. cerevisiae* Hmg2 and human HMGR and a control point between ER-retention and engaging the ER-Golgi trafficking machinery, as in human SCAP and the *S. pombe* analogue Scp1 (13). The involvement of the SSD in a range of pressing human health issues makes it an intriguing target for further studies of regulated misfolding and proteostasis. More broadly, the phenomenon of small-molecule mediated misfolding represents another potential tool for pharmaceutical intervention. In its broadest application, this mallosteric strategy holds untapped potential for targeting proteins less amenable to traditional active site drugs by instead targeting them for either stabilization or for misfolding and degradation.

## Experimental procedures

### Reagents

Geranylgeranyl pyrophosphate (GGPP), cycloheximide, trypsin, and digitonin were purchased from Sigma-Aldrich. Digitonin was washed and recrystallized in ethanol 3 times according to the manufacturer’s instructions. Lovastatin was a gift from Merck & Co (Rahway NJ). GGSPP and 2-fluoro-GGPP were gifts from Reuben Peters (Iowa State University) and Philip Zerbe (University of California Davis). Fos-Choline-13 and Decyl Maltose Neopentyl Glycol were purchased from Anatrace. Anti-myc 9E10 supernatant was prepared from cells (CRL 1729, American Type Culture Collection) cultured in RPMI1640 medium (GIBCO BRL) with 10% fetal calf serum. Living colors mouse anti-GFP monoclonal antibody was purchased from Clontech, and goat anti-mouse antibody HRP-conjugated antibody was purchased from Jackson ImmunoResearch.

### Strains and plasmids

Yeast strains (Supplemental Table 1) and plasmids (Supplemental Table 2) were made by standard techniques. Yeast strains were isogenic and were made from the S288C background. Yeast were grown in rich media (YPD) or in minimal media (Diffco Yeast Nitrogen Base with required amino acids and nucleic acids and 2% glucose) at 30°C.

Hmg2 mutation plasmids were made by splicing by overlap extension (SOEing) or site directed mutagenesis using the Life Technologies GeneArt system.

Ura3 Hmg2-GFP, 1myc_L_-Hmg2-GFP, and Hmg2-1myc plasmids were introduced at the *ura3-52* locus by integration of plasmid cut with *StuI*. Leu2 Hmg2-GFP plasmids were introduced into the promoter of the *leu2Δ* locus by integration of plasmid cut with *PpuMI*.

### Flow cytometry

Flow cytometry experiments were performed as previously described (28). Yeast cultures were grown in minimal media to early log phase. Indicated molecules (22 μM GGPP or 50 μg/mL cycloheximide) or equal volumes of vehicle (7:3 methonol: 10 mM ammonium bicarbonate for GGPP and DMSO for cycloheximide) were added directly to culture medium. Cell fluorescence was measured for 10,000 cells per condition using a BD Accuri C6 flow cytometer (BD Biosciences). Flow data were analyzed using FlowJo software (FlowJo, LLC). Any averages shown are means of 10,000 ungated events.

### Microsome preparation

Microsomes were prepared as described previously (27, 34). Yeast were grown to mid log phase in YPD, and 10 OD equivalents were pelleted, washed in water, and resuspended in 240 μL lysis buffer (0.24 M sorbitol, 1 mM EDTA, 20 mM KH_2_PO_4_/K_2_HPO_4_, pH 7.5) with PIs (2 mM phenylmethylsulfonyl fluoride and 142 mM tosylphenylalanyl chloromethyl ketone). Acid-washed glass beads were added up to the meniscus. Cells were lysed on a multi-vortexer at 4° C for six to eight 1-minute intervals with 1 minute on ice in between each lysis step. The lysates were transferred to a new tube and lysates cleared with 5-second pulses of centrifugation. Microsomes were pelleted from cleared lysates by centrifugation at 14,000 x g for five minutes. Microsome pellets were washed once in XL buffer (1.2 M sorbitol, 5 mM EDTA, 0.1 M KH_2_PO_4_/K_2_HPO_4_, pH 7.5), and resuspended in XL buffer for limited proteolysis.

### Limited proteolysis assay

The limited proteolysis assay was performed as described previously (Shearer & Hampton 2004). Microsomes in XL buffer were treated with the indicated isoprenoid molecules or with equal volumes of vehicle controls. For the S215A kinetic experiments, microsomes were pre-incubated with isoprenoids or vehicle for the indicated times. In all other cases, isoprenoids were added and then samples were immediately incubated at 30° C with 15 μg/mL trypsin. Samples were quenched at the indicated times with equal volumes of 2x urea sample buffer (USB; 8M urea, 4% SDS, 1mM DTT, 125 mM Tris base, pH 6.8) and incubated at 55° C for 10 minutes. Samples were resolved by SDS-PAGE, transferred to nitrocellulose, and blotted with 9E10 anti-myc antibody.

### Detergent limited proteolysis assay

The limited proteolysis assay was performed as described above with the following variations: After initial pelleting and washing of microsomes, microsomes were first thoroughly resuspended in XL buffer and then solubilized by the addition of detergent solution at10x the desired final concentration in XL buffer (final concentration 0.1% Fos-Choline-13, 0.05% DMNG, 0.5% or 1% digitonin). Preparations with detergent were incubated at 4° C for 1 hour with rocking and then repeatedly pipetted up and down. Finally, samples were cleared for by centrifugation in a benchtop microcentrifuge for 15 minutes at 16,000 x g. The supernatants were then separated by ultracentrifugation at 89,000 RPM for 15 minutes, and the supernatant from this step was used for the proteolytic assay. This assay was identical to the limited proteolysis assay described above, except that a lower concentration of trypsin (3 μg/mL instead of 15) was used.

### Detergent solubility assay

Microsomes were prepared as in the detergent limited proteolysis assay up to the point of detergent addition. After thorough mixing, 22 μL of GGPP, 2F-GGPP, GGSPP, or equal volume of vehicle were added as indicated. Microsomes were then incubated at 4° C for 1 hour with rocking. Samples were again pipetted vigorously up and down and then were separated by ultracentrifugation at 89,000 RPM for 15 minutes. Pellet and supernatant were separated and incubated in USB for 10 minutes at 55° C and then resolved and immunoblotted as described above.

## Acknowledgements

The authors wish to thank members of both the Hampton lab and the Pacific Hall 2nd floor laboratory environment for typical vibrant discussions and general good scientific will. We thank Reuben Peters and Philip Zerbe for their generous gift of the isoprenoid GGPP analogs. In addition, RYH would like to express his appreciation to Carbon Hampton for continuous low-key council and tacit approval.

