## Supplemental Table 1 for "An autonomous, but INSIG-modulated, role for the Sterol Sensing Domain in mallostery-regulated ERAD of yeast HMG-CoA reductase"

**Supplemental Table 1. Yeast strains used in this work**

| Strain | Genotype |
| --- | --- |
| RHY3970 | *MATa ade2-101 met2 his3Δ200 lys2-801 leu2Δ ura3-52::URA3::TDH3pr-HMG2-GFP* |
| RHY7661 | MAT*a* *ade2-101 met2 his3Δ200 lys2-801 leu2Δ ura3-52::URA3::TDH3pr-1myc_L_-HMG2-GFP* |
| RHY7683 | MAT*a* *ade2-101 met2 his3Δ200 lys2-801 leu2Δ ura3-52::URA3::TDH3pr-S215A-1myc_L_-HMG2-GFP* |
| RHY11128 | MAT*a* *ade2-101 met2 his3Δ200 lys2-801 leu2Δ ura3-52::URA3::TDH3pr-K357R-1myc_L_-HMG2-GFP* |
| RHY11129 | MAT*a* *ade2-101 met2 his3Δ200 lys2-801 leu2Δ ura3-52::URA3::TDH3pr-K6R-1myc_L_-HMG2-GFP* |
| RHY12225 | *MATalpha ade2-101 met2 his3Δ200 lys2-801 leu2Δ ura3-52::URA3::TDH3pr-1myc_L_-HMG2-GFP YHR133c::kanMX4 YNL156c::KanMX4* |
| RHY12226 | *MATalpha ade2-101 met2 his3Δ200 lys2-801 leu2Δ ura3-52::URA3::TDH3pr-1myc_L_-S215A-HMG2-GFP YHR133c::kanMX4 YNL156c::KanMX4* |
| RHY12239 | *MATa ade2-101 met2 his3Δ200 lys2-801 leu2Δ::TDH3pr-HMG2-GFP ura3-52::URA3::TDH3pr-HMG2-myc* |
| RHY12240 | *MATa ade2-101 met2 his3Δ200 lys2-801 leu2Δ::TDH3pr-HMG2-GFP ura3-52::URA3::TDH3pr-K6R-HMG2-myc* |
| RHY12241 | *MATa ade2-101 met2 his3Δ200 lys2-801 leu2Δ::TDH3pr-HMG2-GFP ura3-52::URA3::TDH3pr-S215A-HMG2-myc* |
| RHY12242 | MAT*a* *ade2-101::ADE2::TDH3pr-Nsg1-3HA met2 his3Δ200 lys2-801 leu2Δ ura3-52::URA3::TDH3pr-1myc_L-_HMG2-GFP* |
| RHY12243 | MAT*a* *ade2-101::ADE2::TDH3pr-Nsg1-3HA met2 his3Δ200 lys2-801 leu2Δ ura3-52::URA3::TDH3pr-S215A-1myc_L-_HMG2-GFP* |
| RHY12244 | MAT*a* *ade2-101 met2 his3Δ200 lys2-801 leu2Δ ura3-52::URA3::TDH3pr-K6R-S215A-1mycHMG2-GFP* |
| RHY12245 | MAT*a* *ade2-101 met2 his3Δ200 lys2-801 leu2Δ ura3-52::URA3::TDH3pr-L219F -1mycHMG2-GFP* |
| RHY12246 | *MATa ade2-101 met2 his3Δ200 lys2-801 leu2Δ::TDH3pr-HMG2-GFP ura3-52::URA3::TDH3pr-1myc_L_-HMG2-GFP* |
| RHY12247 | *MATa ade2-101 met2 his3Δ200 lys2-801 leu2Δ::TDH3pr-HMG2-GFP ura3-52::URA3::TDH3pr-1myc_L_-HMG2-GFP* |
