## Supplemental Table 2 for "An autonomous, but INSIG-modulated, role for the Sterol Sensing Domain in mallostery-regulated ERAD of yeast HMG-CoA reductase"

**Supplemental Table 2. Plasmids used in this work**

| Plasmid | Genotype |
| --- | --- |
| pRH312 | LEU2 YIp |
| pRH313 | URA3 YIp |
| pRH469 | URA3::TDH3pr-HMG2-GFP YIp |
| pRH680 | LEU2::TDH3pr-HMG2-GFP YIp |
| pRH1129 | URA3::TDH3pr-K357R-HMG-GFP YIp |
| pRH1581 | URA3::TDH3pr-1myc_L_-HMG2-GFP YIp |
| pRH1693 | URA3::TDH3pr-K357R-1myc_L_-HMG2-GFP YIp |
| pRH1805 | ADE2::TDH3pr-Nsg1-3HA YIp |
| pRH2177 | URA3::TDH3pr-S215A-1myc_L_-HMG2-GFP YIp |
| pRH2183 | URA3::TDH3pr-L219F-1myc_L_-HMG2-GFP YIp |
| pRH2250 | URA3::TDH3pr-HMG2-1myc |
| pRH2251 | URA3::TDH3pr-K6R-HMG2-1myc |
| pRH2654 | LEU2 TDH3-pr-PGK1term YIP |
| pRH2911 | URA3::TDH3pr-K6R-1myc_L_-HMG2-GFP YIp |
| pRH3210 | URA3::TDH3pr-K6R-S215A-1myc_L_-HMG2-GFP YIp |
| pRH3211 | LEU2::TDH3pr-S215A-HMG2-GFP YIp |
| pRH3212 | URA3::TDH3pr-S215A-HMG2-1myc YIp |
